# Stacked Human Arterial Endothelial Cells Generate Atherosclerotic Fatty Streaks and Release Proinflammatory Cytokines

**DOI:** 10.1101/2024.11.03.621780

**Authors:** Ye Zeng, Zhi Ouyang, Yan Qiu, Wenli Jiang, Chen Jin, Jian Zhong, Linlu Jin, Yixue Qin, Yunran Zhao, Xintong Zhou, Xiaoheng Liu, Bingmei M. Fu

## Abstract

Fatty streaks are the first sign of atherosclerosis. They consist of lipid-containing foam cells, which were believed to be derived from the monocytes in the blood through the leaky endothelium, and from the vascular smooth muscle cells migrating into the intima from the media. Here, we showed that fatty streaks can also be formed by the stacked human arterial endothelial cells (HAECs) cultured in vitro. Via SEM we revealed a novel cell phenotype (coralthelial) that forms a streak/coral-like structure. We observed accumulation of lipids in the coralthelial cell and increased Golgi and coat protein II markers in its nucleus. Additionally, proinflammatory cytokines were upregulated in these cells, likely due to Golgi nuclear translocation and subsequently increased expression of the ribosomal protein RPL23 in the nucleus. We demonstrated, for the first time, that the atherosclerotic fatty streak-like structure can be generated from the stacked HAECs, which also create a proinflammatory microenvironment.

**Significance Statement:** Our work presents a novel perspective on atherosclerotic fatty streak formation, a critical early event in atherosclerosis development. We demonstrate, for the first time, that human arterial endothelial cells (HAECs) can transform into a novel phenotype, termed “coralthelial cells”. These cells contribute to fatty streak formation, and produce proinflammatory cytokines through the relocation of RPL23 with the Golgi apparatus to the nucleus. This finding provides a new cellular mechanism by which endothelial cells initiate atherosclerosis, independent of traditional monocyte-derived foam cells. By enriching established paradigms, our research not only opens new avenues for therapeutic intervention targeting endothelial cells but also enhances understanding of vascular diseases and provides potential diagnostic and prognostic markers for atherosclerosis.

## Introduction

Atherosclerosis is a common condition that develops when a sticky substance (or fatty streak) is formed by fats, cholesterol, and cells inside arterial walls (1, 2). Atherosclerosis-associated cardiovascular diseases are the leading cause of death worldwide. The dysfunction of endothelial cells (ECs) which cover the luminal surface of arterial walls is an initial step during atherosclerosis (3). Endothelial dysfunction or endothelial injury results in increased endothelial permeability, and enhanced proinflammatory cytokines and adhesion molecules that lead to monocyte-endothelial adherence and transmigration. The transmigrated monocytes become activated as macrophages which can be converted to lipid-laden foam cells after engulfing the low-density lipoprotein (LDL) accumulated in the intima through the leaky endothelium (4, 5). The foam cells secrete a variety of cytokines, thereby initiating an inflammatory reaction. In response, vascular smooth muscle cells (VSMCs) in the arterial media migrate into the intima, proliferate, and produce collagen and proteoglycan-rich extracellular matrix (ECM) (6–8). Like macrophages, VSMCs can also turn into fat-filling foam cells via LDL uptake (9). These foam cells construct the fatty streak, and their accumulation is a fundamental pathological factor for the intima thickening (9–11). This conventional view of fatty streak formation and types of cells in the atherosclerotic plaque is depicted in **Fig. 1A**. It was reported that the initial lipid deposition occurred immediately above the internal elastic lamina, rather than near the leaky endothelium, nor in the region dominated by the macrophage foam cells (12–14). The EC-like cells also appeared in the deep intimal regions (9). Lipid accumulation in normocholesterolemic and hypercholesterolemic rabbits following de-endothelialization of the aorta was significantly greater in the re-endothelialized intima than in adjacent regions lacking endothelium. Intimal thickness was enhanced in the re-endothelialized areas for hypercholesterolemic rabbits compared with normocholesterolemic ones, but not in the areas lacking endothelium (11). The thickened intima covered by the regenerated endothelium was named endothelial island where more lipid and lipid-laden foam cells accumulated (**Fig.1A**).

**Figure 1.**
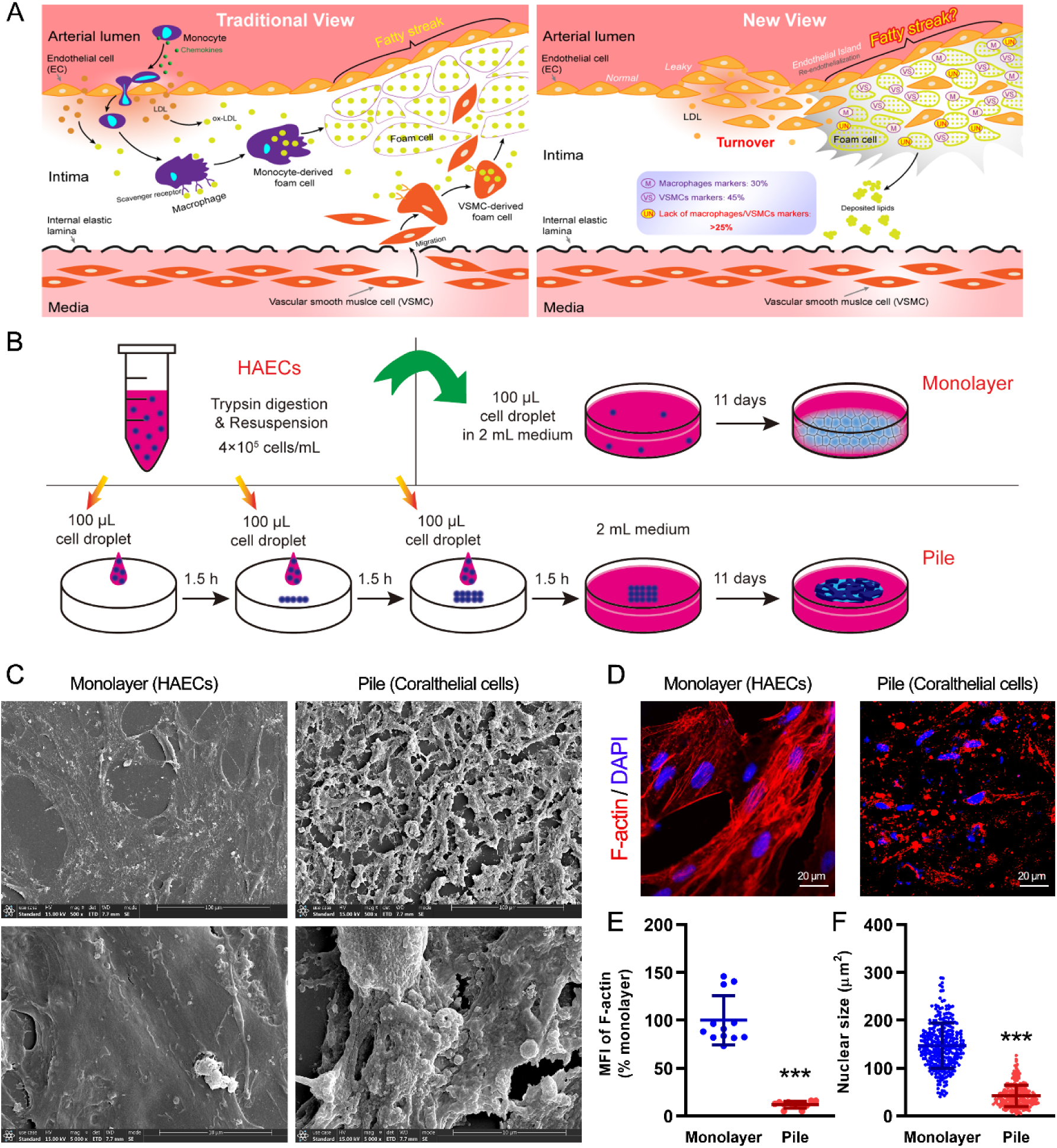
Views on the origin of fatty streaks and their formation by coralthelial cells transformed from the stacked human aortic endothelial cells (HAECs). (A) Schematic of the traditional and new views for the fatty streak. Traditional view (left): Circulating monocytes are attracted by inflammatory chemokines to the sites of the arterial wall with the damaged endothelium. Then they enter the intima via leaky endothelium along with fluid, lipids, and other substances in the blood. The accumulated low-density lipoprotein (LDL) in the intima can undergo oxidative and be avidly taken up by macrophages differentiated from the monocytes and by vascular smooth muscle cells (VSMCs) migrating into the intima from the media to form the monocyte-derived and VSMCs-derived foam cells, respectively. The fatty streak-like structure is constructed by these lipid-containing foam cells in the arterial wall beneath the endothelium. New view (right): The initial lipid deposition occurs in the deep intima close to the internal elastic lamina, rather than near the leaky endothelium, nor in the region dominated by the macrophage foam cells. The intima covered by the regenerated endothelium is significantly thicker and more likely to accumulate lipids. More surprisingly, about 25% or more foam cells in the thickened intima have neither macrophagic markers nor VSMC markers. Based on these observations, we hypothesize that the foam cells without markers for macrophages or VSMCs are derived from endothelial cells (ECs). These ECs come from the rapid turnover after vessel wall damage and are stacked by the local flow in atherosclerosis-prone areas to form an endothelial island. The stacked ECs in the intima can turn into lipid-laden cells and deposit lipids in deep intima along with the lipid-filling foam cells derived from the macrophages and VSMCs. To test this hypothesis, we cultured human arterial endothelial cells (HAECs) after piling them up in the culture dish to simulate the in vivo condition. (B) Schematic of generating monolayer and coralthelial cells (pile) from human aortic endothelial cells (HAECs). After culturing for 11 days, the surface ultrastructure of the monolayer and pile was assessed by scanning electron microscopy (SEM) (C), and the distribution of F-actin cytoskeletons (immuno-stained) and nuclei by confocal microscopy (D). In (C), the bottom panels are close-up views of the top panels. (E) Mean fluorescence intensity (MFI) of F-actin (averaged over 12 fields of 160 μm x 160 μm in each case) and (F) nuclear size (n=300 for each case) were measured from the confocal images. Mean ± standard deviation (SD); ***P<0.001.

While the foam cells in the atherosclerotic intima have been believed to originate from either macrophages or VSMCs (15), only about 30% and 45% of foam cells in atherosclerotic plaques were found to express canonical macrophage and VSMCs markers, respectively (16), suggesting that foam cells have other cell origins. A more recent study by Allahverdian et al (17) showed that 50% of foam cells within advanced human coronary artery lesions expressed the VSMC marker ACTA2, but the majority of these ACTA2+ foam cells also expressed macrophage marker CD68 and represented 40% of all CD68+ lesion cells. By using flow cytometry, Wang et al (18) revealed that 37% of foam cells (BODIPY-labeled cells) were CD45 (monocyte marker)-negative in the atherosclerotic plaque in ApoE-/- mice fed with a normal diet for 27 weeks, but the CD45-negative foam cells were increased to 70% after continued 6 weeks high-fat diet.

The above findings suggest that about 25% or more of the foam cells in the atherosclerotic intima originate from neither monocytes nor VSMCs. We hypothesize that these foam cells are derived from ECs. Experiments by single-cell transcriptome sequencing showed that there were multiple cell clusters in the human atherosclerotic plaque, including ECs, VSMCs and macrophages (19, 20), suggesting that ECs contribute to the foam cell formation in fatty streaks. But where do these ECs come from? Fatty streaks are distributed randomly throughout the arterial tree, however, early lesions are commonly found at sites where changes in blood flow such as a decrease of flow, back currents, or eddy currents occur at branches, bifurcations, and curves in the circulation (10, 21, 22). These observations let us speculate that these ECs come from the rapid turnover after endothelial injury and they are stacked in atherosclerosis-prone areas to form an endothelial island. The stacked ECs in the intima can turn into lipid-laden cells and deposit lipids in deep intima along with the lipid-filling foam cells derived from the macrophages and VSMCs (**Fig. 1A**).

To test this hypothesis, before culturing, we piled human aortic ECs (HAECs) in the culture dish to mimic the packing effect of the disturbed flow in vivo. After 11 days, we employed scanning electron microscopy (SEM) to examine the morphology of the cultured cells after being stacked. To our surprise, instead of a typical cobblestone appearance like HAEC monolayers, these cells showed a coral-like structure. We thus termed them coralthelial cells. We then examined the organelles of the coralthelial cell using transmission electron microscopy (TEM), and lipid accumulation, F-actin cytoskeleton, endothelial markers and coat protein complex II (COPII) via immuno-staining approaches. Moreover, the associated chemokines/cytokines and signaling mechanisms were investigated by RNA sequencing. The secretion of the key identified chemokines/cytokines in the cell culture supernatant was also evaluated by ELISA.

## Results

### The surface ultrastructure and actin cytoskeleton distribution of HAECs (monolayer) and coralthelial cells (pile)

By stacking HAECs before cell culture, a new cell phenotype, coralthelial cells, was formed (**Fig. 1B**). **Figure 1C** shows the surface ultrastructure of the monolayer and the pile (coralthelial cells) revealed by scanning electron microscopy (SEM). After culturing for 11 days, the surface of HAECs in the monolayer appears smooth. A few microvilli, small pits and occasional slender cytoplasmic processes are randomly distributed on the surface of HAECs. In contrast, the culture for the piled cells enabled the three-dimensional growth of HAECs. Stigmata (circular loops) and coral-like structures are observed in the piled HAECs. These piled cells possess a stripe-shape containing a large population of small, spherical, rough blebs on their surface (so named “coralthelial cells”). These cells are morphologically very distinct from the typical flagstone shape of ECs in the monolayer. The cells with the similar coral-like morphology were found in the liver sinusoidal vessel walls, which are thought to be the “fat-storing cells” (23), and also in the focal swelling region of the aorta in the rabbits fed with the cholesterol-rich diet (24).

**Figure 1D** demonstrates the confocal images for the fluorescently immuno-stained F-actin cytoskeleton and nuclei distribution in the HAECs of the monolayer and in the coralthelial cells of the pile. Actin stress fibers in the monolayer are organized in long bundles oriented along the cell extensions or lamellipodia. In contrast, in the pile, the actin fibers are sparse, disorganized, amorphous, and slightly granular in some areas of the cell cytoplasm. In addition, **Figure 1E** shows that the mean fluorescence intensity (MFI) of F-actin in the pile (coralthelial cells) is only 11.75 ± 3.60% of that in the monolayer. These results suggest that the disorganization of the actin meshwork and actin losses contribute to the formation of coralthelial cells. Furthermore, the nuclear size of the coralthelial cells is significantly reduced to 42.16 ± 22.63 μm^2^, compared with 147.10 ± 47.78 μm^2^, the size of the HAECs in the monolayer (**Fig. 1F)**. In coralthelial cells, while glycosaminoglycans (GAGs) were reduced, proteoglycans, glycogen, collagens, and calcium were all enhanced. They can form an ECM scaffold, facilitating the assembly of coralthelial cells into fatty streak-like structure (**Fig. S1**).

### Coralthelial cells have smaller cell bodies, nuclei and mitochondria, but larger autophagic vacuoles and lipid droplets

Figure 2A demonstrates the TEM views of organelles and their quantifications in the HAECs of the monolayer and in the coralthelial cells. In HAECs of the monolayer, there are Weibel-Palade bodies, ribosomes, rough and smooth endoplasmic reticulum (ER), Golgi apparatus and mitochondria in an amorphous ground substance surrounding a nucleus (Fig. 2A). The mitochondria are in the proximity of the rough ER, while Golgi complexes are in the proximity of the smooth ER. There are extensive pinocytic vesicles near the cell junctions and the autophagic vacuoles in the cytoplasm. On the contrary, the morphology and organelles of the coralthelial cells (Fig. 2A) are significantly different from the HAECs in the monolayer. The cell body and nucleus become smaller (**Figs. 2B, C**), as well as mitochondria (Fig. 2D), especially in the horizontal section. The coralthelial cells seem to lack Golgi apparatus and smooth ER in their cytoplasm and the pinocytic vesicles are dramatically reduced near the junctions (Fig. 2A).

**Figure 2.**
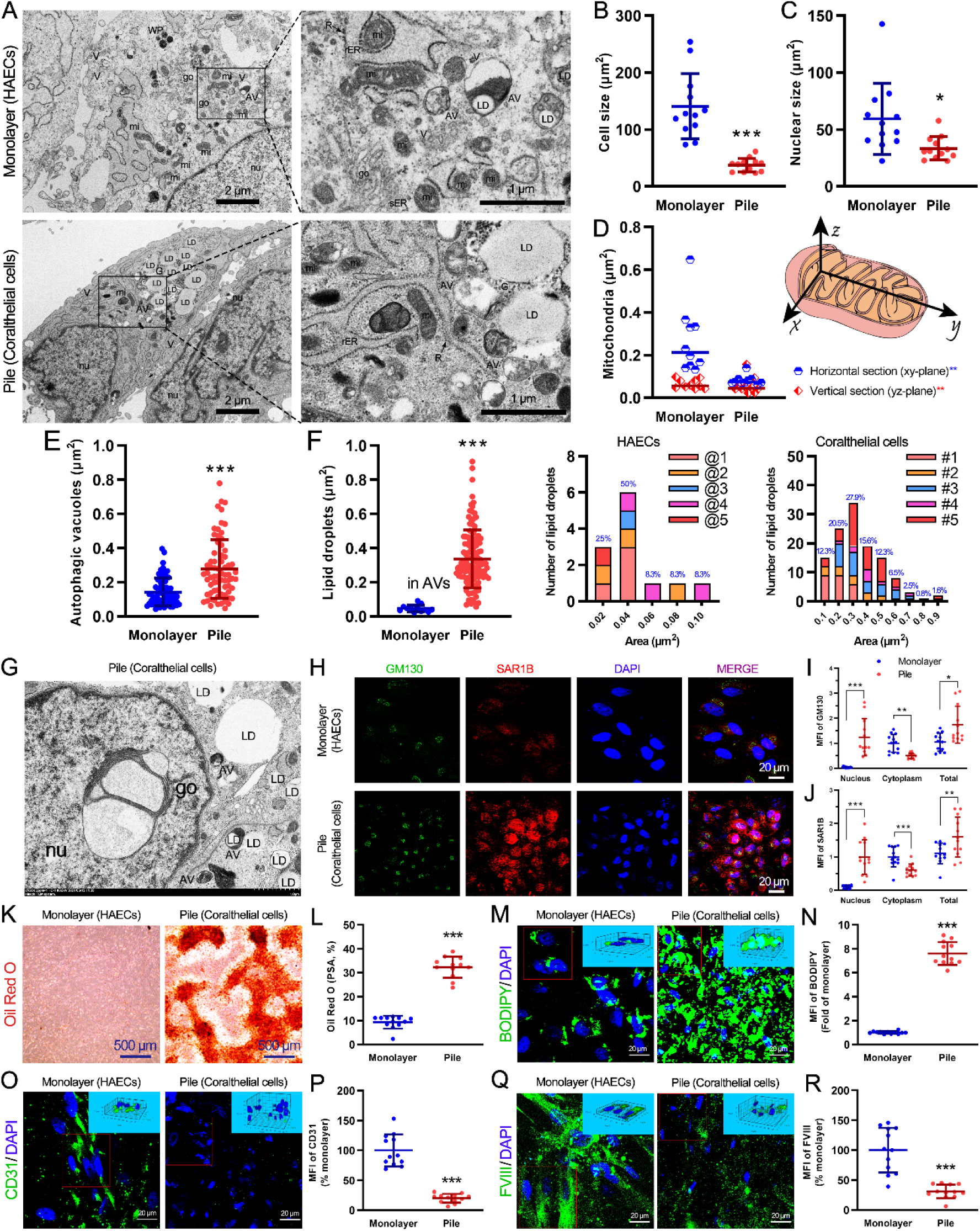
Ultrastructure of coralthelial cells: nuclear import of Golgi and coat protein complex II, lipid deposition, and lack of EC markers. (A) After culturing for 11 days, the ultrastructure and organelles in the cytoplasm were assessed by transmission electron microscopy (TEM) for HAECs in the monolayer and coralthelial cells in the pile. Quantification of the size for cells (B), nuclei (C), mitochondria (D), autophagic vacuoles (E), and lipid droplets (F). (G) The ultrastructure and location of the Golgi apparatus in coralthelial cells. (H) The HAECs from the monolayer and coralthelial cells were co-stained with anti-GOLGA2/GM130 and anti-SAR1B antibodies. The MFIs of GM130 (I) and SAR1B (J) in the nucleus, the cytoplasm, and the total regions per cell were presented as fold of changes relative to the levels in the cytoplasm per HAEC in the monolayer. Staining with Oil Red O (K) and BODIPY lipid probes (M) were performed for the deposition of lipid droplets, (L) and (N) show the corresponding PSA (positive staining area) in percentage %, or MFI (mean fluorescence intensity). They were also stained with endothelial markers, CD31 (O) and FVIII (F8) (Q). (P) and (R) show the corresponding MFI. The 3D views at the right top corner in (M, O, Q) are corresponding to the red boxed regions in these figures. WP: Weibel-Palade body; V: pinocytic vesicles; go: Golgi apparatus; mi: mitochondria; AV: autophagic vacuoles; LD: lipid droplet; G: glycogen; nu: nucleus; R: ribosomes; rER: rough endoplasmic reticulum; sER: smooth endoplasmic reticulum. Mean ± SD; *P<0.05, **P<0.01, ***P<0.001.

The autophagic vacuoles are filled with moderately electrodense flocculent matrixes and lipid droplets. The larger autophagic vacuoles were found in coralthelial cells (**Figs. 2A, E**). Surprisingly, while lipid droplets in the HAECs of the monolayer are few and only appear in autophagic vacuoles, much more and larger lipid droplets were observed in the coralthelial cells (**Figs. 2A, F**). On average, the lipid per cell in the coralthelial cells is ∼10-fold that in the HAECs.

### Translocation of Golgi complexes from cytoplasm to nucleus in coralthelial cells

Since lipids are secreted by smooth ER and Golgi apparatus (25, 26), if there are more lipids in coralthelial cells, there should be more smooth ERs and Golgi apparatus. But coralthelial cells have less Golgi and smooth ER in their cytoplasm compared to HAECs in monolayers. To investigate whether Golgi complexes are translocated from the cytoplasm to the nucleus, we carefully checked the location of Golgi from the TEM images and found Golgi do exist in the nucleus of coralthelial cells (Fig. 2G). We also identified Golgi by using fluorescently conjugated Golgi marker GOGLA2/GM130 (Fig. 2H). Although the amount of GOGLA2/GM130 is less in the cytoplasm of coralthelial cells compared to that of HAECs in the monolayer, it is greatly increased in the nucleus, resulting in more GOGLA2/GM130 per cell in coralthelial cells (Fig. 2I).

Transport of newly synthesized lipids and proteins (26) from ER to Golgi apparatus in eukaryotes is commonly mediated by coat protein complex II (COPII). We thus employed COPII marker SAR1B to identify COPII and found that in consistent with the increased GOGLA2/GM130, SAR1B is also increased in coralthelial cells compared to HAECs in the monolayer (Fig. 2H**-J**). The GM130 and SAR1B are almost absent in the nucleus of HAECs in the monolayer, whereas they are increased in the nucleus and reduced in the cytoplasm of coralthelial cells (Fig. 2H**-J**), suggesting the nuclear translocation of Golgi protein GOGLA2/GM130 by COPII. Colocalization of SAR1B and GOGLA2/GM130 was also observed in the nucleus of coralthelial cells (Fig. 2H).

### Lipid accumulation in coralthelial cells but lack of EC markers

To further verify the lipid accumulation in coralthelial cells, the Oil Red O staining and BODIPY lipid probes were performed (**Figs. 2K-N**). Much more lipid contents were detected in coralthelial cells than HAECs in the monolayer, 32.28 ± 4.43 vs. 9.35 ± 2.64 in Oil Red O positive areas (Fig. 2L). BODIPY-labeled fatty acids appear as a form of agglomeration of irregular sphere particles in coralthelial cells (right figure in Fig. 2M). The MFI of the BODIPY-labeled fatty acids in coralthelial cells is 7.61 ± 0.96 folds of that of HAECs in the monolayer (Fig. 2N).

Two common EC markers CD31 (PECAM-1) and coagulation factor VIII (FVIII, F8) were used to label HAECs in the monolayer and coralthelial cells in the pile (**Figs. 2O and Q**). Interestingly, compared with HAECs in the monolayer, both CD31 and FVIII are significantly decreased in coralthelial cells, 19.95 ± 7.28% in CD31 (Fig. 2P) and 30.97 ± 11.57% in FVIII (Fig. 2R). Although originated from the HAECs, the coralthelial cells are rich in lipids and poor in EC markers, suggesting that they have been transformed from the HAECs to other types of cells, with characters similar to the foam cells in the fatty streak region of the arterial wall in atherosclerosis (27).

### Upregulated inflammation-associated genes and secretion of proinflammatory cytokines in coralthelial cells

To further determine the molecular mechanism that contributes to the generation of coralthelial cells from the HAECs and to screen the associated signaling mechanism, we performed RNA sequencing and identified a total of 339 differentially expressed genes (DEGs) between coralthelial cells and HAECs (Fig. 3A). The principal component analysis (PCA) indicates that the first principal component (PC1: 57.9%) of the DEGs can separate coralthelial cells from HAECs (Fig. 3B). Of all the DEGs, 221 upregulated (in red) and 118 downregulated (in green) in coralthelial cells (Fig. 3A). The proinflammatory cytokines IL-6, MCP-1 and CXCL8 that likely participate causally in human cardiovascular events (28, 29) are among those upregulated genes (in blue) (Fig. 3A).

**Figure 3.**
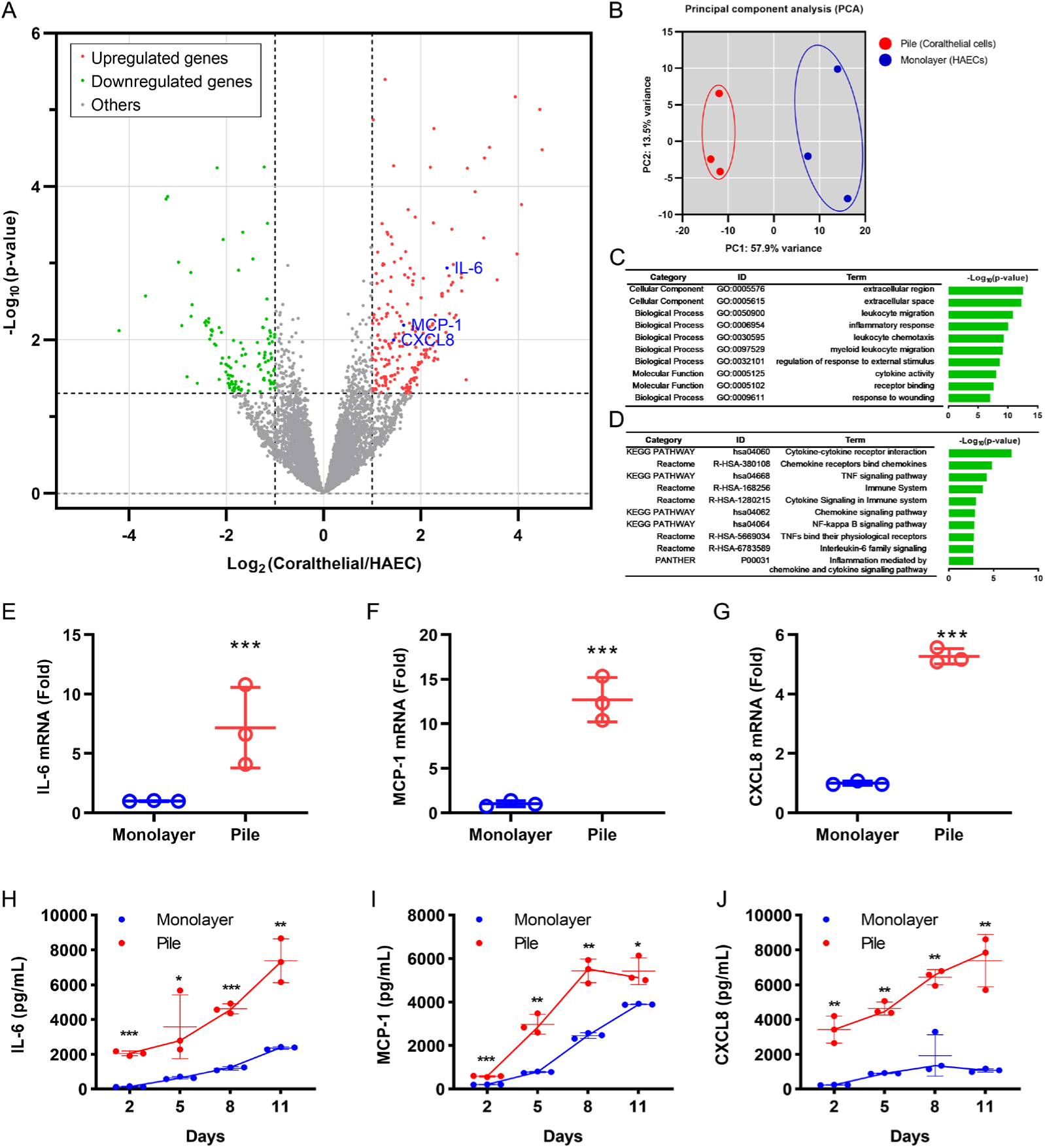
RNA-Seq analysis of coralthelial cells and chemokines/cytokines detection. (A) Volcano plot of DEGs. The red spots indicate the upregulated genes, and the green spots indicate the down-regulated ones. The chemokines/cytokines IL-6, MCP-1 and CXCL8 that have been reported to participate in human cardiovascular activities are labeled in blue. (B) PCA analysis indicates that the transcriptome profile can separate coralthelial cells (pile) from HAECs (monolayer) with the first principal component (PC1). (C) GO analysis of DEGs indicates that GO terms related to chemokines/cytokines in inflammation are significantly altered in coralthelial cells. (D) Pathway analysis of DEGs indicates that chemokines/cytokines signaling pathways in inflammation are also significantly altered in coralthelial cells. (E-G) qRT-PCR confirms the increased mRNA expression of IL-6 (E), MCP-1 (F), and CXCL8 (G) in coralthelial cells. (H-J) ELISA detection of IL-6 (H), MCP-1 (I), and CXCL8 (J) in the culture medium at day 2,5,8,11 for coralthelial cells and HAECs. *P<0.05; **P<0.01; ***P<0.001 vs. HAECs (monolayer).

By Gene Ontology (GO) analysis, the top enriched cellular component items for all DEGs are “extracellular region” and “extracellular space”, the top enriched biological process items are “leukocyte migration”, “inflammatory response”, “leukocyte chemotaxis”, “myeloid leukocyte migration”, “regulation of response to external stimulus”, and “response to wounding”; and the top enriched molecular function items are “cytokine activity” and “receptor binding” (Fig. 3C). These results indicate that the chemotaxis and inflammatory response are altered in coralthelial cells. The results of the pathway analysis are consistent with the GO analysis. The top enriched pathways are 4 KEGG terms: cytokine-cytokine receptor interaction, TNF signaling pathway, chemokine signaling pathway and NF-κB signaling pathway; 5 Reactome terms: chemokine receptors bind chemokines, immune system, cytokine signaling in immune system, TNFs bind their physiological receptors, and Interleukin-6 (IL-6) family signaling; and 1 PANHER term: inflammation mediated by chemokine/cytokine signaling pathway (Fig. 3D). These enriched pathways suggest that coralthelial cells play a critical role in inflammatory chemokine/cytokine activity.

Using qRT-PCR, we showed that the mRNA levels of the key proinflammatory cytokines IL-6, MCP-1 and CXCL8 in coralthelial cells are increased to 7.18 ± 3.40 (Fig. 3E), 12.71 ± 2.49 (Fig. 3F), and 5.28 ± 0.26 (Fig. 3G) folds of that in HAECs, respectively. We further confirmed the secretion of these cytokines by coralthelial cells via ELISA. In the culture supernatant, IL-6, MCP-1 and CXCL8 of coralthelial cells are markedly increased with the culturing time, and are significantly higher than those of monolayer HAECs from day 2 to day 11 (Fig. 3H**-J**). For example, at day 11, 7378.71 ± 1256.92 vs. 2365.56 ± 69.86 pg/mL for IL-6; 5426.25 ± 617.89 vs. 3897.84 ± 20.93 pg/mL for MCP-1; and 7385.77 ± 1504.66 vs. 1079.0 ± 92.72 pg/mL for CXCL8, respectively, in the supernatant of coralthelial cells vs. that in monolayer HAECs.

### Roles of Golgi complex nuclear translocation and RPL23 in the generation of proinflammatory cytokines

The ribosomal protein RPL23 is closely associated with the production of proinflmmatory mediators in ECs and intima growth in response to disturbed flows (30, 31). To investigate the role of RPL23 in creating a proinflammation microenvironment, we labeled RPL23 (red), as well as fibrillarin (FBL, green), which is specifically expressed in the nucleoli. The nucleolar structure was disrupted and became irregular in the coralthelial cells compared to that of HAECs in the monolayer (Fig. 4A). There was a remarkable increase in the localization of RPL23 in the nucleolus of coralthelial cells (red overlapped with the green), suggesting that RPL23 increased in the nucleolus, accompanied by the increased nucleolar stress, as indicated by the redistribution of the nucleolar stress responses indicator fibrillarin (FBL) (Fig. 4A).

**Figure 4.**
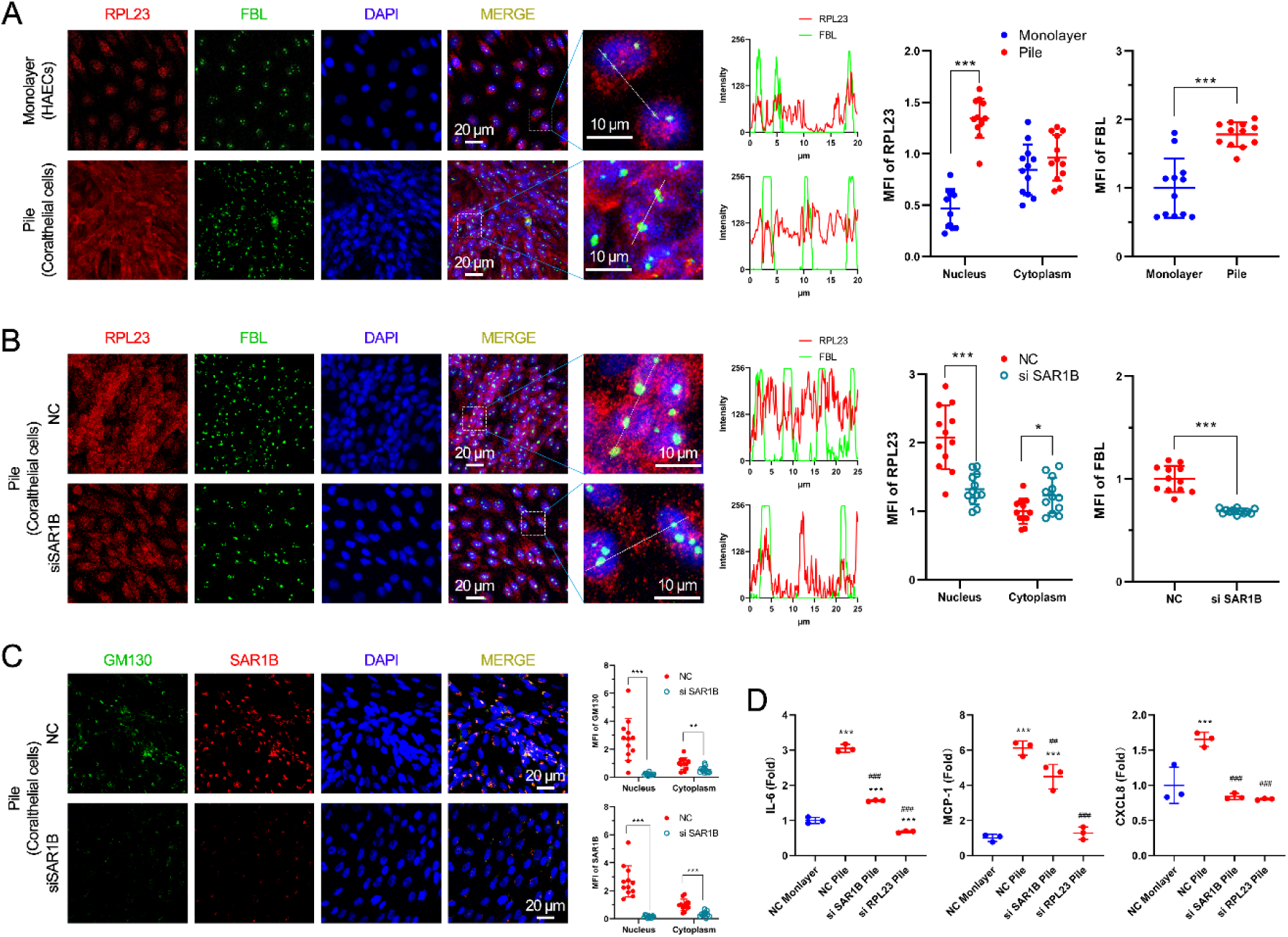
RPL23 translocates into the nucleolus and facilitates the secretion of proinflammatory cytokines, which is attenuated by SAR1B siRNAs. (A) Co-staining of RPL23 and FBL in HAEC monolayer and coralthelial cells. (B-C) Following SAR1B knockdown, the nuclear translocation of RPL23 (B) and GM130 (C) were assessed in coralthelial cells. Lines scan for RPL23 and FBL were performed. The MFIs of RPL23, FBL, GM130 and SAR1B in the nucleus and cytoplasm are presented as fold changes relative to their levels in the cytoplasm of HAEC in monolayer or NC coralthelial cells (pile). Mean ± SD; *P < 0.05, **P < 0.01, ***P < 0.001. (D) The levels of IL-6, MCP-1 and CXCL8 in coralthelial cells were quantified. ***P<0.001 vs. NC monolayer; ##P<0.01, ###P<0.01 vs. NC pile.

Protein trafficking between the ER and the Golgi apparatus is essential for cellular processes, including the release of proinflammatory cytokines. To further explore the impact of nuclear translocation of COPII vesicles and Golgi apparatus in the coralthelial cells, we attempted to suppress COPII by knocking down SAR1B using siSAR1B. Strikingly, siSAR1B did not restore the nucleolar structure but inhibited RPL23 translocation to the nucleolus (Fig. 4B). This suggests that siSAR1B may disrupt intranuclear transport in the Golgi apparatus, considering that transport from the ER to the Golgi is typically mediated by COPII. siSAR1B not only suppressed the expression of SAR1B but also reduced GM130 and its nuclear translocation (Fig. 4C), further suggesting that RPL23 is translocated into the nucleus together with GM130. RPL23 is used for 60S subunit assembly (32), and is located at the exit tunnel of the ribosome, serving as a docking site for the nascent polypeptide-associated complex (NAC) chaperone in protein folding (33). Inhibition of Golgi nuclear translocation results in reduced RPL23 in the nucleus and, consequently, less RPL23 in the nucleolus, possibly leading to decrease in newly translated proteins.

To investigate the requirement of RPL23 for the synthesis of proinflammatory cytokines, we examined the secretion of key proinflammatory cytokines including IL-6, MCP-1, and CXCL8 in the coralthelial cells transfected with siSAR1B and siRPL23. The secretion of these cytokines by coralthelial cells was significantly suppressed by both siSAR1B and siRPL23 (Fig. 4D). Overall, these findings suggest that coralthelial cells secrete chemokines/cytokines to create a proinflammatory microenvironment through the trafficking of RPL23 with Golgi apparatus in the nucleus.

## Discussion

The fatty streak is the initial sign of atherosclerosis which causes many clinical complications, including coronary heart disease, stroke, and peripheral vascular disease. It is widely known that the fatty streaks in the arterial intima are formed by the lipid-laden foam cells derived from the macrophages differentiated from the circulating monocytes via the leaky endothelium and from the vascular smooth muscle cells (VSMCs) migrating into the intima from the media. In the present study, we presented compelling evidence that human arterial endothelial cells (HAECs) can directly transform into fat-laden foam cells after stacking them up in culture, without the need for atherogenic stimuli. These transformed cells, which have a coral-like appearance, are called “coralthelial cells”. Similarly shaped cells have been observed in the liver sinusoidal vessel wall, which were thought to be the “fat-storing cells” (23). Such cells are also present in the focal swellings in the aorta in the early stage of atherosclerosis (24).

Through TEM and lipid-specific staining techniques, including Oil Red O staining and BODIPY lipid probes, we observed numerous lipid droplets inside the coralthelial cells. Unlike the foam cells derived from monocytes or VSMCs, which tend to be larger, coralthelial cells are much smaller than normal HAECs, about 25% in the cell size and 50% in the nucleus size. To our surprise, these coralthelial cells lack typical vascular EC markers CD31 and FVIII, suggesting that they may evade detection within atherosclerotic plaques or be misclassified as monocyte-or VSMC-derived foam cells due to their phenotypic transformation. In addition, the cytoskeleton of coralthelial cells appears disorganized, and fatty acids within the cells aggregate into irregular spherical particles, contrasting with the small, uniformly distributed spherical vesicles observed in the VSMCs incubated with oxidized low-density lipoprotein (ox-LDL) (34). These distinct cytological features suggest that coralthelial cells represent a novel cellular phenotype potentially involved in lipid deposition within the atherosclerotic plaques. However, further research is needed to identify specific phenotype markers for coralthelial cells.

Both macrophages and VSMCs are known to internalize natural LDLs. However, the internalization of natural LDL does not cause intracellular cholesterol accumulation or foam cell formation. For a long time, ox-LDL has been thought to be the primary form of modified LDL taken up by macrophages and VSMCs to result in foam cell formation during atherosclerosis (35). Many in vitro studies have employed high concentrations of ox-LDL (50-100 μg/mL) to stimulate monocytes or VSMCs for 12-72 h to induce lipid droplets biogenesis and foam cell formation (36–38). Although ox-LDLs are highly cytotoxic to ECs, vascular ECs can take up the active component of ox-LDL lysophosphatidylcholine (1 μg/mL for 24 h) by phagocytosis (39). Hypercholesterolemic serum can also transform the mouse aortic ECs into foam cells, and disorganize the cytoskeleton (40). In contrast to adding atherogenic agents to induce foam cells, we simply cultured the HAECs after stacking them up. Foam cell formation is typically driven by an imbalance in cholesterol influx, esterification, and efflux (41). The overexpression of the scavenger receptor LOX-1 in endothelial cells has been shown to induce the endothelial dysfunction and exacerbate atherosclerosis in the apoE^-/-^ mice (42). Such metabolic dysfunctions may also contribute to the transformation of coralthelial cells, promoting lipid accumulation and subsequent fatty streak formation.

A critical question is whether the in vitro induction of fat-filling coralthelial cells reflects in vivo processes. By scanning electron microscope (SEM), it has been observed an increased incidence of lipid-rich fatty dots and streaks in the aortic arch of rabbits fed an atherogenic diet (43). The early lesions appear as focal swelling surrounded by morphologically normal endothelium and possibly covered by rounded ECs. Crystal-like structures and small focal swellings have been observed on the luminal surface of aortas in these rabbits. Numerous focal swellings are hypothesized to comprise the early fatty streaks (43). Analyses of fresh human atherosclerotic samples have indeed shown that the presence of lipid and lipid-crystalline droplets within the fatty streaks (44). These droplets are primarily composed of cholesteryl esters and triglycerides, covered with a surface of phospholipids and unesterified (free) cholesterol in the human aorta (45). We speculate that in vivo, HAECs undergo rapid turnover following endothelial injury and are piled up, possibly due to the vascular spasms and local disturbed flow commonly observed at the branches, bifurcations, and curves in the vascular system where fatty freaks are initiated.

The recruitment of circulating monocytes and T cells to atherosclerotic sites is driven by the proinflammatory cytokines. The coralthelial cells secret proinflammatory cytokines, including IL-6, MCP-1, and CXCL8, which are well-documented contributors to atherosclerosis progression. For instance, IL-6 recruits T cells to sites of inflammation, while IL-8 and MCP-1 serve as chemotactic agents for leukocytes (46, 47). RNA sequencing revealed significant upregulation of inflammatory pathways in the coralthelial cells, especially those related to cytokine-cytokine receptor interactions, TNF signaling, and NF-κB signaling. These pathways likely promote leukocyte migration and exacerbate local inflammation, contributing to the sustained proinflammatory microenvironment observed in atherosclerotic plaques.

A pivotal finding of this study is the nucleolar translocation of the RPL23 ribosomal protein, which is essential for the production of proinflammatory cytokines. RPL23 encodes a cytoplasmic ribosomal protein that serves as a component of the 60S subunit (32). Elevated RPL23 expression is associated with a reduction in apoptosis in the CD34+ bone marrow cells via c-MYC upregulation (48). The nucleolus, a critical stress sensor and signaling hub, responds early to cellular damage as nucleolar stress. Impaired incorporation of RPL23 into maturing 60S subunits enhances nucleolar stress responses under oncogenic stress (32). In triple-negative breast cancer cells, translocation of RPL23 to the nucleolus triggers c-MYC expression and promotes cellular stemness (49). Similarly, in the coralthelial cells, we observed nucleolar stress and increased RPL23 accumulation. SAR1B is a GTP-binding protein involved in the assembly of the COPII vesicle and the transport of lipids and proteins from the ER to the Golgi apparatus (26). Knockdown of SAR1B and RPL23 significantly reduced cytokine secretion, indicating that RPL23 plays a crucial role in the ER-Golgi-nucleus axis, mediating inflammation in the coralthelial cells. RPL23 has also been implicated in intimal growth in response to disturbed flow (30, 31), linking mechanical forces, endothelial cell stacking, and proinflammatory microenvironment during atherosclerosis.

Finally, our study highlights the importance of Golgi-ER dynamics in cellular transformation and inflammation. GOGLA2/GM130 and SAR1B were previously thought to localize exclusively in the cytoplasm of normal endothelial cells (26, 50). To our surprise, we did observe their unexpected translocation to the nucleus in the coralthelial cells, alongside RPL23. These findings suggest that COPII vesicles and Golgi apparatus, normally restricted to the cytoplasm, translocate to the nucleus in the coralthelial cells. Inhibition of the Golgi apparatus, particularly via SAR1B knockdown, disrupted RPL23 nuclear translocation and cytokine secretion, underscoring the therapeutic potential of targeting the Golgi-ER axis in the treatment of atherosclerosis.

In conclusion, we demonstrated, for the first time, that the lipid-laden cells (coralthelial cells) can be directly derived from HAECs without the need for atherogenic factors. These cells not only contribute to lipid deposition but also create a proinflammatory microenvironment through Golgi nuclear translocation and RPL23 expression in the nucleolus (Fig. 5), potentially exacerbating plaque formation and disease progression. Our findings present a novel paradigm for understanding atherosclerosis. Future research should investigate the mechanisms driving the transformation of HAECs into coralthelial cells and explore the potential of targeting Golgi-ER trafficking pathways as a therapeutic strategy for mitigating inflammation and plaque development in atherosclerosis. Furthermore, identifying specific markers for coralthelial cells would facilitate their detection and classification within atherosclerotic plaques, allowing for a more accurate understanding of their roles in disease progression. Finally, elucidating the interplay between mechanical forces, endothelial cell turnover, and coralthelial cell formation could provide deeper insights into how disturbed flow contributes to the initiation and propagation of fatty streaks and plaques. These investigations hold promise for novel therapeutic approaches aimed at reducing the burden of cardiovascular diseases associated with atherosclerosis.

**Figure 5.**
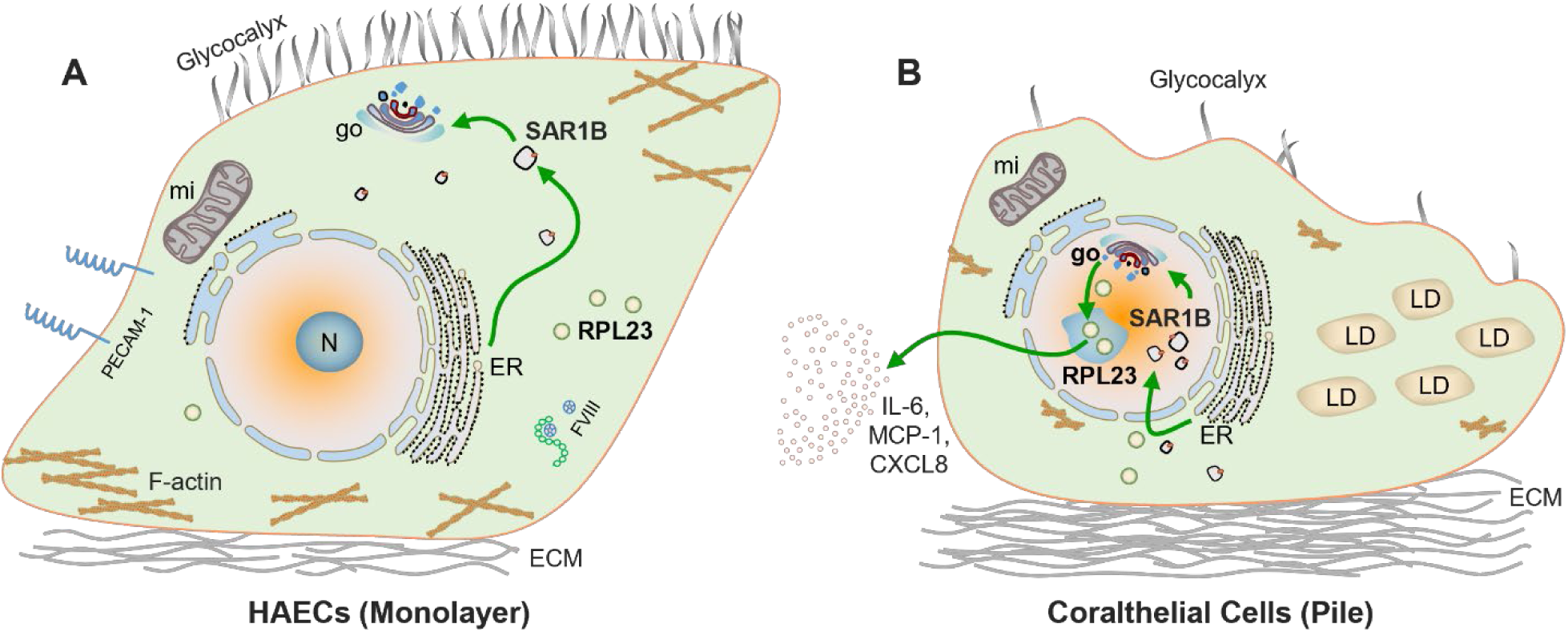
The major differences between HAECs and coralthelial cells, as well as the mechanism of cytokine release in coralthelial cells. (A) HAECs display a typical endothelial morphology with a flattened, elongated structure, organized actin filaments, intact glycocalyx, and minimal lipid accumulation. (B) In contrast, coralthelial cells, derived from stacked HAECs, exhibit a distinct coral-like morphology with surface blebbing, smaller cell bodies and nuclei, disorganized actin filaments, degraded glycocalyx, smaller mitochondria, enhanced nucleolar stress, and significant lipid droplet accumulation, looking like foam cells in atherosclerotic plaques. Coralthelial cells also lack typical endothelial markers such as PECAM-1 (CD31) and FVIII, which are present in HAECs. Coralthelial cells show a marked increase in collagen production, contributing to the formation of the extracellular matrix (ECM) scaffold observed in atherosclerotic fatty streak-like structures. In addition, coralthelial cells show a significant upregulation of proinflammatory cytokines, including IL-6, MCP-1 and CXCL8, which promote a proinflammatory microenvironment characteristic of atherosclerosis. This cytokine release in coralthelial cells is likely driven by the ER-Golgi-nucleus axis. It involves the translocation of RPL23 from the cytoplasm to the nucleus, in conjunction with the Golgi apparatus and COPII vesicles (key component SAR1B), which initiates cytokine production. Targeting RPL23 and ER-Golgi dynamics presents a promising therapeutic strategy for modulating cytokine release in vascular diseases, including atherosclerosis. go: Golgi apparatus; ER: endoplasmic reticulum; mi: mitochondria; LD: lipid droplet; N: nucleolus.

## Materials and Methods

Detailed methods are described in *SI Materials and Methods, Fig. S2, and Table S1*.

## Acknowledgments

This project was supported by grants from: the National Natural Science Foundation of China 12272246 (YZ) and 11932014 (XHL), the National Institutes of Health UG3UH3TR002151 (BMF), and the Key Research and Development Projects of Sichuan Province 2023YFS0075 (YZ).

## Supporting Information

## Materials and Methods

### Cell culture

Human aortic endothelial cells (HAECs; Sciencell, USA) were cultured in a humidified 5% CO_2_ incubator at 37°C. After confluence, the cells were trypsinized with trypsin-EDTA (0.05%) (#15400054, Invitrogen, USA), and then concentrated by centrifugation at x 176 g for 5 min (3H16RI, HereXi Instrument & Equipment, Hunan, China). The harvested cells were resuspended with fresh endothelial cell medium (ECM, #1001, Sciencell, USA) and adjusted to a concentration of 4×10^5^ cells/mL.

TeloHAECs (#CRL-4052, ATCC, USA) were used for siRNA transfection and to investigate the relationship between RPL23 and Golgi nuclear translocation, nucleolar stress, and were maintained under the same conditions as HAECs. To knock down the SAR1B and RPL23, the siRNAs were used and the transfection was performed using Opti-MEM and lipofectamine 3000 according to the manufacturer’s instructions. After 48 h, cells were used for further analysis. The knockdown efficiency was determined by evaluating the levels of SAR1B and RPL23 mRNA expression using qPCR (**Fig. S2**). **Table. S1** details the sequences of the primers used.

### Induction of coralthelial cells

For the induction of coralthelial cells, 100 μL cell suspension was firstly dropped in the center of a 35 mm culture dish (#430165, Corning, USA), and then placed in the incubator for 1.5 h. After nonadherent cells were removed, another 100 μL cell suspension was dropped at the original position and incubated for 1.5 h. After nonadherent cells were removed the second time, a third 100 μL cell suspension was added and incubated for 1.5h. Finally, 2 mL fresh medium was added to the culture dish after removing nonadherent cells and cultured for 11 days at 37°C in a 5% CO_2_ incubator.

The HAECs cultivated into a monolayer were used as the control. A total of 100 μL cell suspension was mixed with 2 mL fresh medium and cultured for 11 days at 37°C in a 5% CO_2_ incubator to form a monolayer.

### Scanning electron microscopy (SEM)

To evaluate cell surface morphology, the HAECs in the monolayer and the coralthelial cells in the pile were prepared for SEM. Briefly, the samples were firstly washed with phosphate buffer solution (PBS) (#BL302A, Biosharp, China), fixed overnight in 3% glutaraldehyde (#G810415, Macklin, China) in 0.1M PBS (pH7.4). After sequential dehydration in ethanol (30%, 50%, 70%, 80%, 90%, 100% and 100%) for 15 min each, the cell samples were dried in a critical point drier (EM CPD300, Leica, Vienna, Austria) for 1.5 h, sputtered with gold (MC1000, Hitachi, Tokyo, Japan) and viewed under field emission SEM (Helios G5 UC, Thermoisher Scientific, USA) equipped with an EDS (AZteclive Ultim Max 100, Oxford Instruments, UK) at 15kV.

### Cytoskeleton assessment

Polymerized actin filaments (F-actin) were evaluated by phalloidin staining. Cells were fixed for 10 min with 4% paraformaldehyde (#P0099, Beyotime, China), washed three times in PBS, incubated with 0.1% Triton X-100 (#ST795, Beyotime, China) for 5 min and Actin-Tracker Red-555 (1:200, #C2203S, Beyotime, China) for 60 min at room temperature followed by three washes with PBS. Finally, cells were stained with DAPI (#C1005, Beyotime, China) for 5 min and then observed by the confocal laser scanning microscopy. As previously described (1), all fluorescent staining samples were imaged by a confocal laser scanning microscope (LSM 710, Carl Zeiss, Germany) using a Plan-Apochromat 63×/1.4 oil DIC objective. The quantification analyses were performed using ImageJ software (version 1.53q; National Institutes of Health, USA).

### Transmission electron microscope (TEM)

Cells were firstly fixed in 3% glutaraldehyde in 0.1 M PBS (pH 7.4) at 4°C for 12 h and post-fixed in 1% osmium tetroxide in 0.1 M PBS at 4°C for 2 h. After fully rinsing in distilled water, cells were dehydrated in graded acetone series (30%, 50%, 70%, 90%, 100%, and 100%, each for 10 min) and embedded in SPI-Pon 812R epoxy resin (#02659R-AB, SPI, USA). The above steps were completed on the glass slide and then cells were separated from the slide. Ultrathin sections (70 nm) were cut using an ultramicrotome (EM UC7, Leica, Austria), and attached to copper grids (AN200, Japan) with Formvar film (Beijing Zhongjingkeyi Technology Co., Ltd., China), stained with 2% uranyl acetate (#02624-AB, SPI) and lead citrate (#02532-BA, SPI) and imaged by TEM (Hitachi HT7800, Tokyo, Japan) at 80 kV.

### Oil Red O staining and quantification

The lipid accumulation in cells was detected by Oil Red O staining. Cells were firstly washed three times with PBS and fixed in 4% paraformaldehyde for 10 min followed by three washes with deionized water. Oil Red O (#G1262, Solarbio, China) was thoroughly mixed with distilled water in a ratio of 3:2 and filtered with neutral filter paper (Whatman, USA). After staining with Oil Red O for 30 min, cells were rinsed with deionized water and observed under an inverted phase-contrast microscope (EVOS XL Core, Thermo Fisher Scientific, USA). For quantification, the dye of Oil Red O was extracted by 100% isopropanol for 20 min, and the absorbance at 510 nm was measured by a full wavelength micrometer (SpectraMax 190, Molecular Devices, USA).

### BODIPY fluorescent probe

A BODIPY probe was used to detect the cellular lipid droplets. Cells were firstly fixed in 4% paraformaldehyde for 10 min, washed in PBS three times for 5 min each, and then stained with 2 μM BODIPY 493/503 (#D3922, Molecular Probes, USA) for 15 min, along with DAPI (3 nM; #D1306, Invitrogen, USA) counterstaining for 5 min. Then, cells were observed by the confocal laser scanning microscopy.

### Canonical EC makers immunofluorescence staining

Cells were fixed for 10 min with 4% paraformaldehyde followed by three washes in PBS, permeabilized with 0.1% Triton X-100 for 5 min, incubated with 1% BSA (#A7284, Sigma Aldrich, USA) in PBS for 20 min, and then incubated with rabbit anti-CD31 (1:200, #CSB-PA017767LA01HU, CUSABIO, China) and anti-F8 (FVIII) antibodies (1:200, #CSB-PA007932LA01HU, CUSABIO, China) for 1 h. After three washes in PBS, cells were incubated with Alexa 488 conjugated goat anti-rabbit IgG (1: 400, #CA11034s, Invitrogen, USA) at room temperature for 1 h and DAPI (Invitrogen, USA) for 5 min followed by three washes in PBS, and finally observed by the confocal laser scanning microscopy.

### Detection of glycocalyx by wheat germ agglutinin (WGA) staining

Wheat germ agglutinin (WGA) binds to sialic acid and N-acetyl-D glucosamine of the glycocalyx. WGA staining has been used to detect and quantify the glycocalyx of cells. Cells were incubated with fluorescein isothiocyanate-labeled WGA (#L4895, FITC-WGA; Sigma Aldrich, USA) at 5 μg/mL for 1 h at room temperature in the dark. After washing in PBS three times for 5 min each, slides were counterstained with DAPI (Invitrogen, USA) for 5 min and observed under the confocal laser scanning microscopy. The orthogonal view of the WGA stack was performed using Zen blue 3.6 software (Zeiss, Germany). The quantification analyses were performed using ImageJ software.

### Toluidine Blue staining

The sulfate and/or carboxylic groups in mucopolysaccharides (acidic glycosaminoglycans) and proteoglycan complexes were stained by Toluidine Blue staining (#G3660, Solarbio, China). Cells were washed three times with PBS, fixed with 95% ethanol for 15 s, rinsed with deionized water, stained with 500 μL Toluidine Blue solution for 5 min and maintained in deionized water for 15 min. After rinsing with deionized water, the samples were observed under an inverted phase contrast microscope.

### Periodic Acid-Schiff (PAS) staining

A PAS staining kit (#G1360, Solarbio, China) was used to detect the carbohydrate aggregation. Cells were washed three times with PBS, fixed with the PAS fixative (Reagent A) for 15 min, washed three times with deionized water and air-dry, and treated with the oxidant (Reagent B) at room temperature for 15 min. Then, cells were rinsed with deionized water, stained in the Schiff reagent (Reagent C) at room temperature for 20 min in dark, rinsed with the sodium sulfite solution (Reagent D) twice for 2 min each, and then redyed with the Mayer Hematoxylin staining solution for 2 min, rinsed with deionized water and observed.

### Picrosirius Red Staining

Picrosirius Red staining (#S8060, Solarbio, China) was performed for collagen detection. Cells were washed three times with PBS, fixed in 4% paraformaldehyde for 10 min, washed three times with deionized water, stained with 0.1% Sirius Red solution for 30 min, rinsed with deionized water and observed.

### Alizarin red S staining

Alizarin red S staining (#G1452, Solarbio, China) was performed to detect the calcium deposition. Cells were washed three times with PBS, fixed in 4% paraformaldehyde for 10 min, washed three times with PBS, stained with 0.2% Alizarin Red S solution for 20 min, rinsed with PBS and observed.

### Locations of Golgi, coat protein complex II (COPII), RPL23, and fibrillarin (FBL) by immunofluorescence staining

Cells were fixed for 10 min with 4% paraformaldehyde followed by three washes in PBS, permeabilized with 0.1% Triton X-100 for 5 min. Nonspecific binding sites were blocked via incubating in 3% BSA in PBS for 30 min, and then were incubated with mouse anti-GOLGA2/GM130 antibodies (1:200, #AG2041, Beyotime, China) and rabbit anti-SAR1B (1:200, #22292-1-AP, Proteintech, USA), or rabbit anti-RPL23 (1:200, #A4292, ABclonal, China) and mouse anti-FBL (1:200, #66985-1-Ig, Proteintech, USA) for 1 h at room temperature, followed by washing three times in PBS and subsequent incubation with Alexa Fluor 488 conjugated goat anti-mouse IgG (1:400, # CA11001s, Invitrogen, USA) and Alexa Fluor 555 conjugated donkey anti-Rabbit IgG (1:400, #A0453, Beyotime, China) at room temperature for 1 h. The samples were further stained with DAPI (Invitrogen, USA) for 5 min. Finally, samples were washed in PBS and were imaged using the confocal laser scanning microscope. The quantification analyses were performed using ImageJ software.

### RNA sequencing and identification of chemokines/cytokines

RNA from the HAECs and coralthelial cells was extracted using Trizol reagent (#15596-0180, Life Technologies, USA) according to the manufacturer’s instruction, followed by integrity evaluation, mRNA-seq libraries constructing, and high-throughput sequencing using an Illumina Genome Analyzer (CapitalBio Corp, Beijing, China). The mRNA sequencing raw reads were filtered firstly by removing the low-quality reads, filtering the contaminants, and trimming the adaptor sequences. The unigenes resulting from the assembly of the mRNA reads were annotated, and the expression levels of the mRNA transcripts were measured as previously described (2). The differentially expressed genes (DEGs) were determined with p-value < 0.05 and the absolute log_2_-fold change not less than 1.0. Gene ontology (GO) and pathway enrichment analysis were performed. DEGs in the enriched GO terms and pathways related to chemokines and cytokines in inflammation (p-value < 0.05) were identified using the KEGG Orthology Based Annotation System (KOBAS 3.0, http://kobas.cbi.pku.edu.cn/kobas3).

### Principal component analysis (PCA)

Principal component analysis (PCA) was performed using GraphPad Prism 9 (Version 9.2.0, GrahPad Software, USA) to assess the gene expression patterns in the coralthelial cells and HAECs.

### Quantitative real-time qRT-PCR

RNA was extracted from the cells using TRIzol (Invitrogen, USA), cDNA synthesis using PrimeScript™ RT reagent Kit with gDNA Eraser (#RR047A, Takara, Japan) and an oligo (dT) primer. qRT-PCR assays were performed using a 2×Taq SYBR Green qPCR Premix (SQ101-01, Innovagene, China) on Applied Biosystems QuantStudio^TM^1 Real-Time PCR System (Thermo Fisher Scientific, USA). qRT-PCR was performed in 10 μL of quantitative reaction mixtures containing 5 μL of SYBR Premix Ex Taq (#RR390A, Takara, Japan), 0.4 μM each of forward and reverse primers, 1 μL cDNA, and nuclease-free water. The program used for qRT-PCR consisted of a denaturing cycle of 3 min at 94 °C, 40 cycles of PCR (94 °C for 10 s, 60 °C for 40 s) and a melting cycle of 60 °C for 15 s. The primer sequences are listed as below: IL-6, 5’-GCC TTC GGT CCA GTTGCCTTC-3’ (forward) and 5’-GTTCTGAAGAGGTGAGTGGCTGTC-3’ (reverse); CXCL8, 5’-CTC TCT TGG CAGCCTTCCTGATTTC-3’ (forward) and 5’-GGGGTGGAAAGGTTTGGAGTATGTC-3’ (reverse); MCP-1(CCL2): 5’-ACCAGCAGCAAGTGTCCCAAAG-3’ (forward) and 5’-TTTGCTTGTCCAGGTGGTCCATG-3’ (reverse); SAR1B, 5’-GGGTGGACATGTTCA AGCTCGA-3’ (forward) and 5’-TCGCAACCTCTCTTCACTGATGG-3’ (reverse); RPL23, 5’-GGTGATGGCCACAG TCAAGA-3’ (forward), 5’-CGTTGTCGAATGA CCACTGC-3’ (reverse), and GAPDH, 5’-TGTTCGTCATGGGTGTGAAC-3’ (forward) and 5’-ATGGCATGGACTGTGGTCAT-3’ (reverse).

The mRNA levels of IL-6, CXCL8, and MCP-1 were normalized to the GAPDH mRNA level. Results of real-time PCR were analyzed using the 2^−ΔΔCT^ method to compare the transcriptional levels of each gene in coralthelial cells to that in HAECs.

### Enzyme-linked immunosorbent assay (ELISA)

The level of IL-6, MCP-1, and CXCL8 in the culture supernatant of HAECs and coralthelial cells at 2, 5, 8, and 11 days were detected by human IL-6 ELISA Kit (#ELH-IL6, RayBio, USA), MCP-1 ELISA Kit (#ELH-MCP1, RayBio), and CXCL-8 ELISA Kit (#ELH-IL8, RayBio) in accordance with the manufacturer’s instructions.

### Statistical analysis

Student’s *t* tests or one-way ANOVA were used to determine statistical significance between groups using SPSS software (v26; IBM, USA). *P* < 0.05 was considered statistically significant.

In addition to lipid and foam cells, collagens and ECM contribute to the intimal thickening (3). Collagens comprise a major portion of proteins in the ECM found in the atherosclerotic plaque. In the lesion-free intima from 23 human subjects, out of 100 mg dry matter, 2.5 mg is mucopolysaccharide, 25 mg is collagen, and 9.8 mg is lipid. In contrast, in the fatty streak region, there is an increase of 0.33 mg in mucopolysaccharide, paralleled by an increase of 3.3 mg in collagen but 22 mg in lipid per 100 mg dry tissue (4). VSMCs are thought to be the main cell type responsible for collagen synthesis in the vessel wall. VSMCs orchestrate the assembly of type I collagen fibril that is linked to the cytoskeleton (5). Meanwhile, type I collagens are usually present in the diseased vessels and support the proliferation of VSMCs (6).

### Reduced glycosaminoglycans (GAGs) and enhanced proteoglycans in coralthelial cells

The endothelium is intact over large lesions. It is believed that the ECs overlying the lesion are abnormally large while they lose argyrophilic properties with unclear cell boundaries (7). Loss of silver precipitation at the lesion areas is thought to be associated with the degradation of glycocalyx during atherosclerotic lesion development (7, 8). Glycocalyx plays a critical role in vascular homeostasis (9), and its coverage and thickness are reduced even before the atherosclerotic plaque formation (10). At plaque sites, particularly at bifurcations, glycocalyx is markedly diminished on ECs (11). The degradation of the glycocalyx is closely associated with increased artery stiffness caused by hypertension and aging (12, 13). It should be noted, however, that an early report demonstrated that the concanavalin A reactive coat (glycocalyx) over endothelium varies in thickness during lesion development. Initially, it becomes thicker (0.2-0.6 µm), but eventually becomes amorphous depositions adhering to the denuded endothelial surface (14).

The glycocalyx at the cells was examined by staining with wheat germ agglutinin (WGA) and toluidine blue (**Figs. S1A-D**). WGA specifically binds to N-acetyl-D-glucosamine and N-acetyl-D-neuraminic acid of GAGs (15). Toluidine blue is a cationic dye with high affinity with the sulfate and/or carboxylic groups of mucopolysaccharides (acid GAGs) and proteoglycan complexes (16). While WGA-stained GAGs are distributed on the surface of HAECs in the monolayer (left two panels in **Fig. S1A**), GAGs are largely reduced in the coralthelial cells and dispersed in their cytoplasm (right two panels in **Fig. S1A**). The MFI of GAGs at the coralthelial cells is only 32.77 ± 8.85% of HAECs in the monolayer (**Fig. S1B**). However, the extracellular matrix proteoglycan complexes are significantly increased in the coralthelial cells (**Fig. S1C**), compared with that in the HAECs of the monolayer, 17.95 ± 1.48% vs. 7.06 ± 1.56% (**Fig. S1D**). The residual proteoglycans are predominantly distributed in the coral-like structures.

### Enriched glycogen, collagens, and calcium in coralthelial cells

The accumulated periodic acid Schiff (PAS) positive materials in the coralthelial cells are significantly increased compared to those in the HAECs of the monolayer, 32.53 ± 2.96% vs. 7.79 ± 0.76% (**Figs. S1E, F**), along with much more toluidine blue positive materials (**Figs. S1C and D**), supporting that the glycogen and residual proteoglycans deposits are aggregated in the coral-like structures. Picrosirius red staining was performed to quantify collagen deposition (**Fig. S1G**). Coralthelial cells appear to have more intense staining of collagen, 27.47 ± 2.75% vs. 4.62 ± 2.41%, compared to HAECs in the monolayer (**Fig. S1H**). Collagen fibrils appear to scaffold the coral-like structures. Calcium and mineralization within cells were assessed by Alizarin red staining (**Fig. S1I**). Substantially more Alizarin red staining was seen in coralthelial cells than that in HAECs of the monolayer, 23.35 ± 4.13% vs. 2.37 ± 0.55% (**Fig. S1J**). Like proteoglycans, glycogen and collagen, calcium is also distributed in the coral-like structures of coralthelial cells. It appears that the coralthelial cells (or transformed HAECs) can synthesize collagen fibrils as a supporting network for the accumulation of lipid droplets, proteoglycans, glycogens and calcium to form the fatty streak-like structure.

Our findings suggest that HAEC-derived coralthelial cells also produce much more collagens than original HAECs. The collagens form an ECM scaffold, facilitating the assembly of coralthelial cells into fatty streak-like structure. Proteoglycans, glycogen, and calcium were also abundant in these structures. Proteoglycans (17), collagen type I (18) and sulfated glycosaminoglycans (19), which enhance intimal retention of plasma LDLs, probably enter the autophagic vacuoles to mix with droplets. We also observed autophagic vacuoles filled with moderately electrodense flocculent matrixes and lipid droplets in the coralthelial cells.

---

**Fig. S1.**
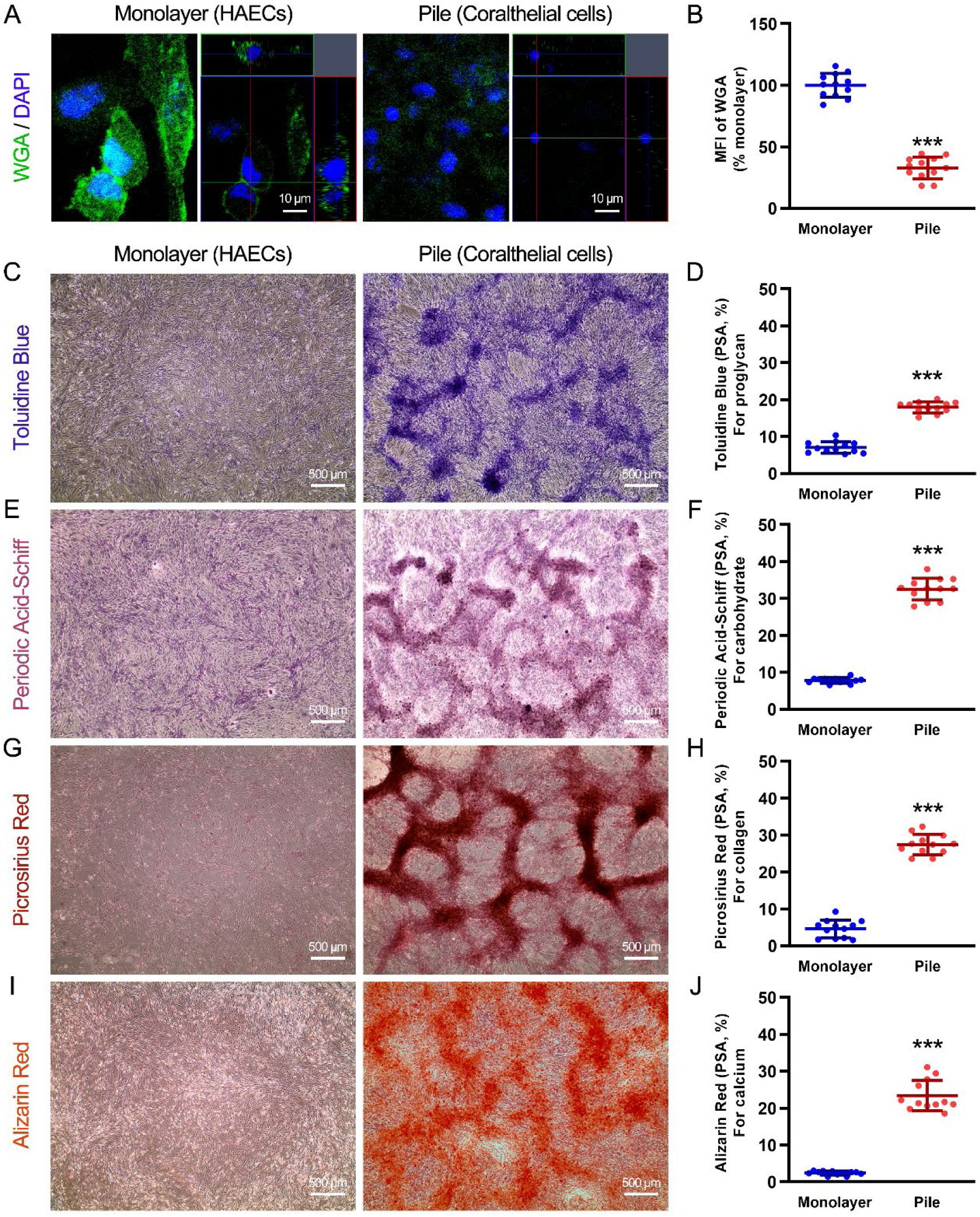
Distribution of glycocalyx, glycogen, collagens and calcium in HAECs in the monolayer and coralthelial cells in the pile. After culturing for 11 days, HAECs from the monolayer and coralthelial cells from the pile were stained with WGA for glycosaminoglycans (**A**), toluidine blue for mucopolysaccharides and proteoglycan complexes (**C**), Periodic Acid-Schiff (PAS) for carbohydrate aggregation (**E**), Picrosirius red for collagen (**G**), and Alizarin red for calcium and mineralization (**I**). The corresponding mean fluorescence intensity (MFI) of these stains is shown in (**B, D, F, H, J**), respectively. Mean ± SD; **P<0.01, ***P<0.001.

**Fig. S2.**
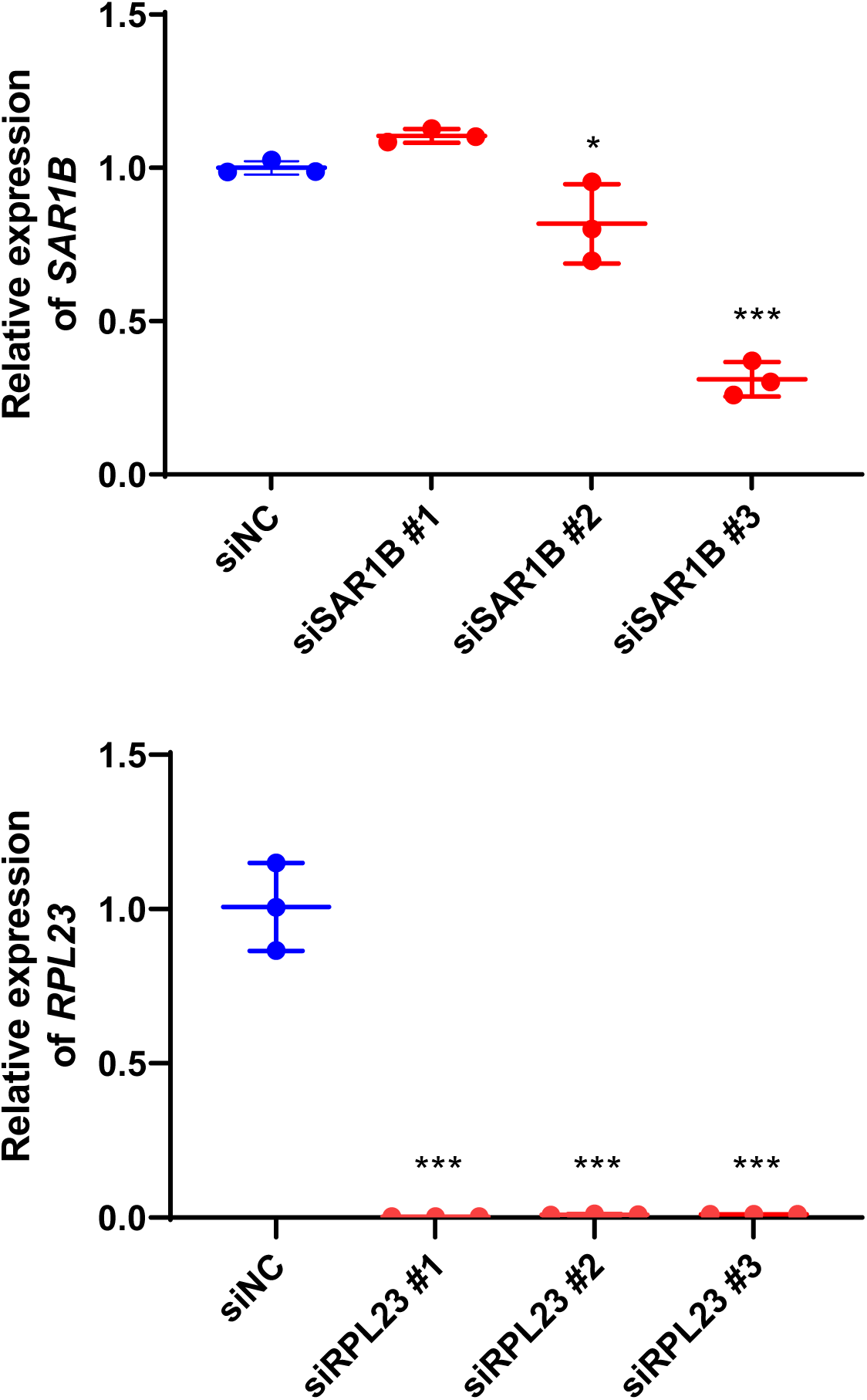
The interference efficiency of the small interfering RNAs for SAR1B and RPL23. SAR1B-3 and RPL23-3 were used in following experiments. Mean ± standard deviation (SD); * P<0.05, ***P<0.001 vs. siNC.

**Table S1.**
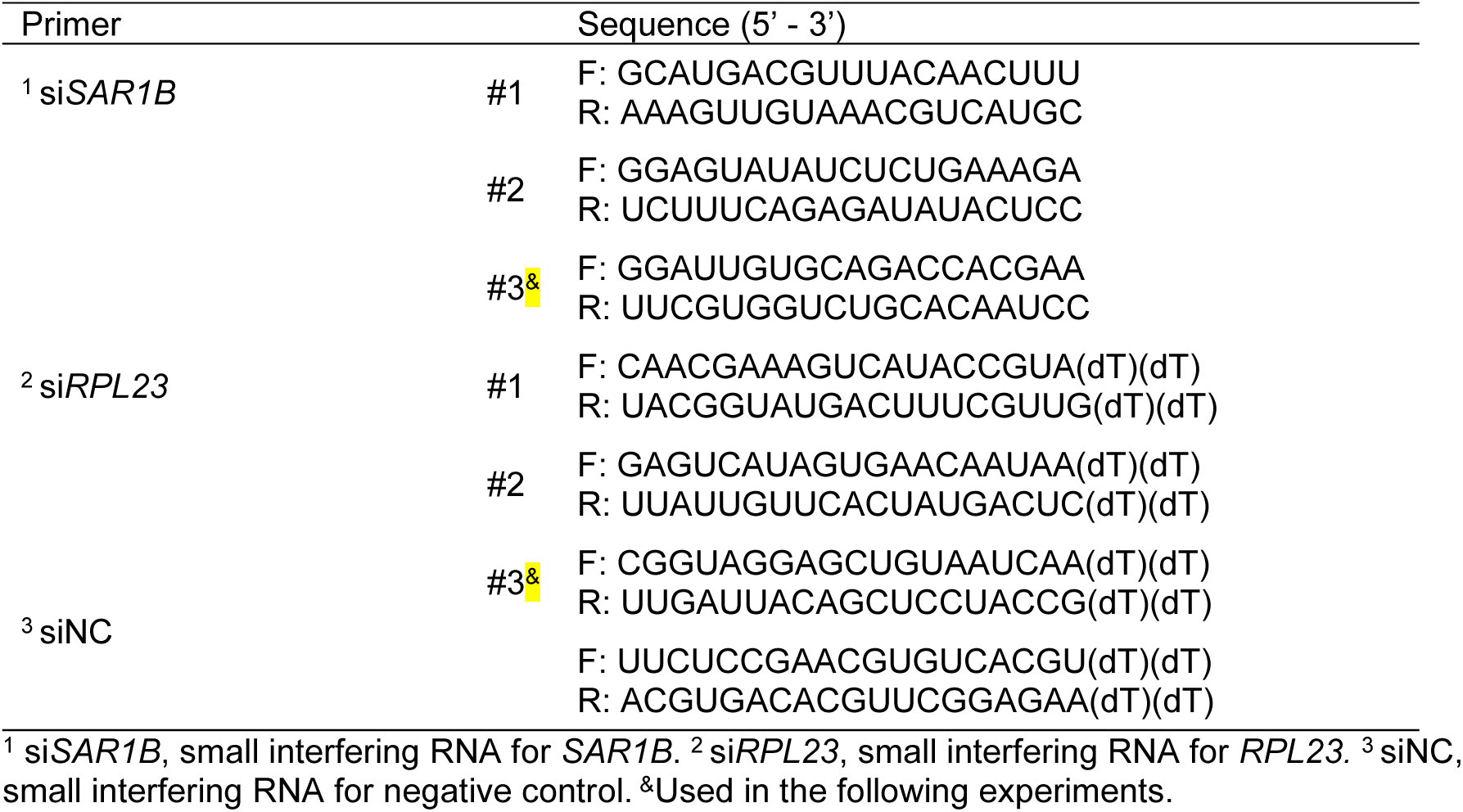
siRNA sequences for RNA interference.

